# Galvanic current activates the NLRP3 inflammasome to promote type I collagen production in tendon

**DOI:** 10.1101/2021.09.20.461048

**Authors:** Alejandro Peñín-Franch, José Antonio García-Vidal, Carlos Manuel Martínez, Pilar Escolar-Reina, Ana I.Gómez, Francisco Minaya-Muñoz, Fermín Valera-Garrido, Francesc Medina-Mirapeix, Pablo Pelegrín

## Abstract

The NLRP3 inflammasome coordinates inflammation in response to different pathogen- and damage-associated molecular patterns, being implicated in different infectious, chronic inflammatory, metabolic and degenerative diseases. In chronic tendinopathies lesions, different non-resolving mechanisms produce a degenerative condition that impairs tissue healing, complicating their clinical management. Percutaneous needle electrolysis consist in the application of a galvanic current and is emerging as a novel treatment for tendinopathies. Here we found that galvanic current activates the NLRP3 inflammasome and the induction of an inflammatory response promoting a collagen-mediated regeneration of the tendon. This study establish the molecular mechanism of percutaneous electrolysis for the treatment of chronic lesions and the beneficial effects of an induced inflammasome-related response.

## INTRODUCTION

Tissue damage and infection triggers an inflammatory response that is coordinated by the activation of the immune system. Inflammation promotes a response to recover homeostasis by removing invading pathogens and repairing tissues (Medzhitov, 2008). However, chronic inflammation can induce the continuous production of tissue regenerative factors and an excessive accumulation of components of the extracellular matrix leading to tissue fibrosis (Alegre et al., 2017; Borthwick et al., 2013; Gaul et al., 2020; Wynn, 2008). Therefore, an equilibrated inflammatory response is required to recover homeostasis and successfully achieve tissue healing (Borthwick et al., 2013; Eming et al., 2017; Liston and Masters, 2017; Medzhitov, 2008). In response to tissue damage, the nucleotide-binding oligomerization domain with leucine rich repeat and pyrin domain containing 3 (NLRP3) inflammasome is activated and coordinates an inflammatory response (Broz and Dixit, 2016; Schroder et al., 2011). NLRP3 inflammasome is a multiprotein complex mainly formed in myeloid cells after encounter damage- or pathogen-associated molecular patterns, including elevated concentrations of extracellular ATP, changes in extracellular osmolarity or detection of insoluble particles and crystals, as uric acid crystals or amyloid deposition (Amores-Iniesta et al., 2017; Compan et al., 2012; Heneka et al., 2013; Mayor et al., 2006). NLRP3 oligomers recruit the accessory apoptosis-speck like protein with a caspase recruitment and activation domain (ASC) that favor the activation of the inflammatory caspase-1 (Boucher et al., 2018; Li et al., 2018; Schmidt et al., 2016). Caspase-1 proteolytically process immature pro-inflammatory cytokines of the interleukin (IL)-1 family to produce the bioactive form of IL-1β and IL-18 (Broz and Dixit, 2016; Schroder et al., 2011). Caspase-1 also process gasdermin D protein (GSDMD), and its amino-terminal fragment (GSDMD^NT^) oligomerize in the plasma membrane forming pores allowing the release of IL-1β and IL-18 cytokines, as well as other intracellular content, including inflammasome oligomers (Baroja-Mazo et al., 2014; Broz et al., 2020).

NLRP3 activation occurs in different chronic inflammatory, metabolic and degenerative diseases such as gout, type 2 diabetes or Alzheimer (Daniels et al., 2016; Heneka et al., 2013; Masters et al., 2010; Mayor et al., 2006), therefore selective small molecules that block NLRP3 are emerging as novel anti-inflammatory therapies (Cocco et al., 2017; Coll et al., 2015; Tapia-Abellán et al., 2019). However, in some pathological circumstances, a boost, rather than an inhibition of NLRP3 would be beneficial to reduce clinical complications, such as in immunosuppressed septic patients that accumulate high mortality rates due to secondary infections associated to a profound deactivation of the NLRP3 inflammasome (Martínez-García et al., 2019). In chronic non-resolving lesions, as tendinopathies developed after prolonged extreme exercise, there are several mechanisms establishing a degenerative condition of the tissue that impairs healing and complicate clinical management (Cook and Purdam, 2009; Regan et al., 1992; Soslowsky et al., n.d.). Anti-inflammatory therapies have shown inefficient in randomized trials for the treating of this type of lesions (Bisset et al., 2006; Coombes et al., 2013) and novel treatments are emerging aiming at the regeneration of the tissue (Bubnov, 2013; Chellini et al., 2019), including the minimally invasive percutaneous needle electrolysis (De-la-Cruz-Torres et al., 2020; Margalef et al., 2020; Valera-Garrido et al., 2013, 2014, 2020). Percutaneous needle electrolysis consist in the application of a galvanic current through an acupuncture needle, combining mechanical and electrical stimulation in the tissues, resulting in a local controlled microtrauma that derives in an inflammatory response and the repairment of the affected tissue (Valera-Garrido and Minaya-Muñoz, 2019). However, the detailed molecular mechanism behind percutaneous needle electrolysis inducing the induction of an inflammatory response has not been yet described. In this study, we found that galvanic current applicated during percutaneous needle electrolysis was able to activate the NLRP3 inflammasome and induce the release of IL-1β from macrophages. Mice deficient on NLRP3 failed to increase IL-1β in tendons after percutaneous needle electrolysis and resulted in a reduction of TGF-β and type I collagen deposition, indicating that the NLRP3 inflammasome plays an important role in the regenerative response of the tendon associated to percutaneous needle electrolysis.

## RESULTS

### Galvanic current enhances macrophage pro-inflammatory M1 phenotype

We initially designed and produced a device to apply galvanic current to adherent cultured cells in 6 well cell culture plates (**Fig. S1**), this device allowed us to explore the effect of galvanic currents in bone marrow derived mouse macrophages. Application of 2 impacts of 12 mA of galvanic current for 6 seconds each over LPS stimulated macrophages, induced an increase of the expression of *Cox2* and *Il6* genes (**Fig. 1A**). However, it did not affect LPS-induced *Il1b* or *Tnfa* pro-inflammatory gene expression (**Fig. 1A**). Interestingly meanwhile *Tnfa* expression was upregulated with galvanic current alone (**Fig. 1A**), galvanic currents were not inducing the expression of *Cox2, Il6* or *Il1b* genes on non-LPS treated macrophages, or over IL-4 treated macrophages (**Fig. 1A**). When macrophages were polarized to M2 by IL-4, galvanic currents decreased the expression of the M2 markers *Arg1, Fizz1* and *Mrc1* (**Fig. 1B**). These data suggest that galvanic current could enhance the pro-inflammatory signature of M1 macrophages whilst decrease M2 polarization. We next studied the concentration of released pro-inflammatory cytokines from macrophages, and found that galvanic current was not able to increase the concentration of IL-6 or TNF-α release after LPS stimulation (**Fig. 1C**), but significantly augmented the release of IL-1β in an intensity dependent manner (**Fig. 1C**). This data indicates that the increase of *Il6* and *Tnfa* gene expression detected at mRNA level would not be transcribing to higher amounts of released IL-6 and TNF-α over LPS treatment, but galvanic current could be potentially activating an inflammasome to induce the release of IL-1β.

**Figure 1.**
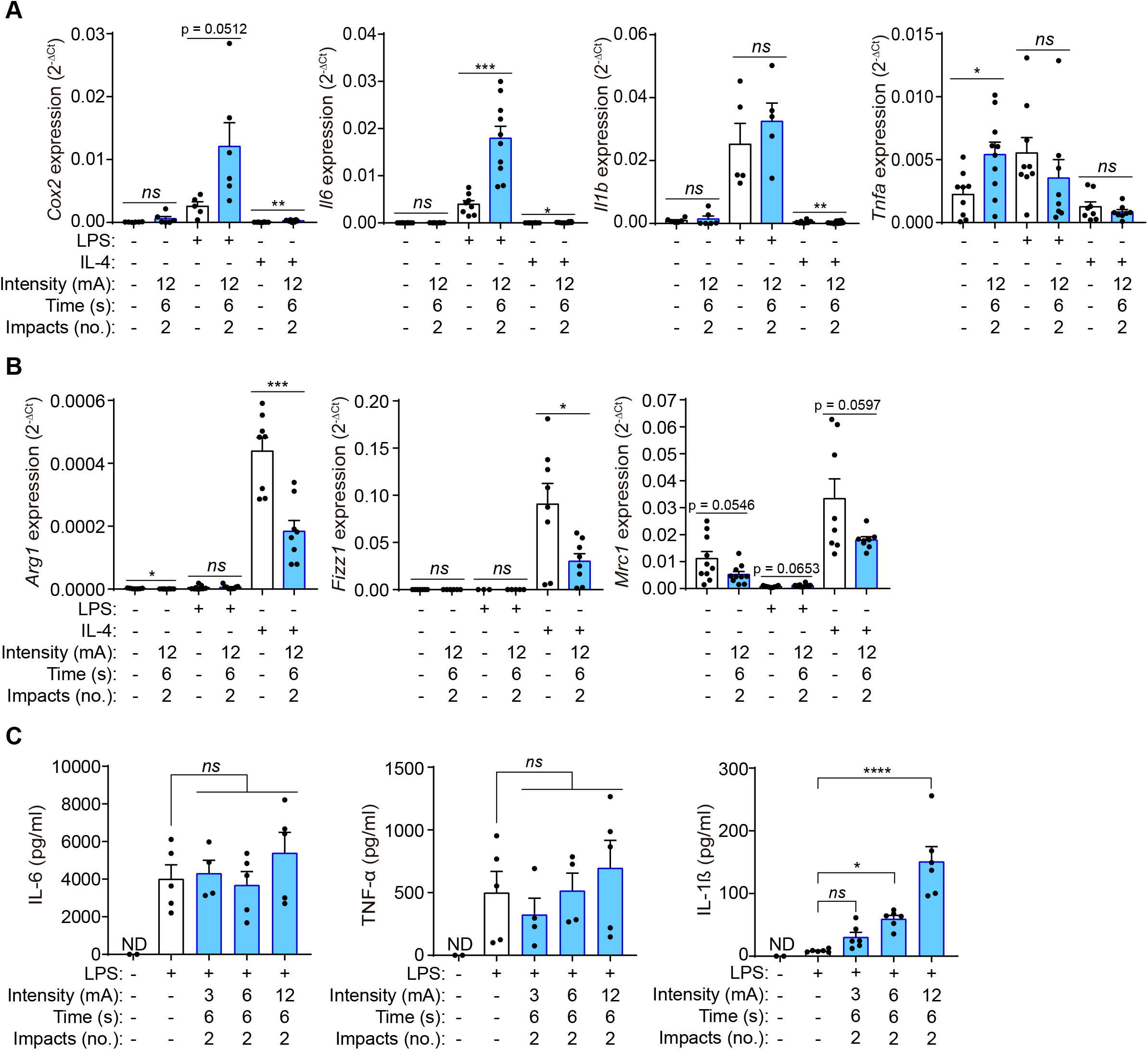
Galvanic current increases the M1 phenotype of macrophages. (**A**) Quantitative PCR for M1 genes *Cox2, Il6, Il1b* and *Tnfa* expression from mouse bone marrow derived macrophages (BMDMs) treated for 2 h with LPS (1 μg/ml) or 4 h with IL-4 (20 ng/µl) as indicated and then 2 impacts of 12 mA of galvanic current for 6 seconds was applicated. Cells were then further cultured for 6 h before analysis. Center values represent the mean and error bars represent s.e.m.; *n*= 5-10 samples of 5 independent experiments; for *Cox2* unpaired *t*-test except in the two first colums where Mann-Whitney test were performed, for *Il6* Mann-Whitney test except in LPS *vs* LPS+galvanic current comparison where unpaired *t*-test were performed, for *Il1b* Mann-Whitney test, for *Tnfa* unpaired *t*-test except in LPS *vs* LPS+galvanic current comparison where Mann-Whitney were performed, \*\*\**p*<0.0005, \*\**p*<0.005, \**p*<0.05, and *ns p*>0.05. (**B**) Quantitative PCR for M2 genes *Arg1, Fizz1* and *Mrc1* expression from BMDMs treated as in (A). Center values represent the mean and error bars represent s.e.m.; *n*= 3-10 samples of 5 independent experiments; unpaired *t*-test except LPS *vs* LPS+galvanic current comparison where Mann-Whitney test were performed, \*\*\**p*<0.0005, \**p*<0.05, and *ns p*>0.05. (**C**) IL-6, TNF-α and IL-1β release from BMDMs treated as in (A) but different intensities of galvanic current (3, 6, 12 mA) were applicated. Center values represent the mean and error bars represent s.e.m.; *n*= 2 for untreated cells and *n*= 4-6 for treatment groups from 4 independent experiments; one-way ANOVA were performed comparing treated groups with control group, \*\*\*\**p*<0.0001, \**p*<0.05, and *ns p*>0.05.

### Galvanic current activates the NLRP3 inflammasome

Since IL-1β release is increased by the activation of caspase-1 after the canonical or non-canonical inflammasome formation (Broz and Dixit, 2016), we next studied the release of IL-1β induced by galvanic current in macrophages deficient on caspase-1 and -11 to avoid both the canonical and non-canonical inflammasome signaling. We found that *Casp1/11*^-/-^ macrophages fail to release IL-1β induced by galvanic current (**Fig. 2A**). We then found that galvanic current application on *Pycard*^-/-^ macrophages also failed to induce the release of IL-1β, denoting that the inflammasome adaptor protein ASC would be also required for the inflammasome activation (**Fig. 2A**). Since current application could be considered a sterile danger signal, we next assessed the implication of NLRP3, an inflammasome sensor important to elicit an immune response in sterile dangerous situations (Broz and Dixit, 2016; Liston and Masters, 2017). *Nlrp3*^-/-^ and the use of the specific NLRP3 inhibitor MCC950 (Coll et al., 2015; Tapia-Abellán et al., 2019) impaired the release of IL-1β induced by galvanic current (**Fig. 2A,B**), demonstrating that the NLRP3 inflammasome is activated during galvanic current application. As controls, similar results were obtained in parallel with the specific NLRP3 activator nigericin (**Fig. 2B and S2A**). Mechanistically, the use of an extracellular buffer with 40 mM of KCl decreased IL-1β release induced by nigericin and galvanic current application, but not the release of IL-1β induced by *Clostridium difficile* toxin B, that activate the Pyrin inflammasome which is a K^+^-efflux independent inflammasome (**Fig. 2C**). However, meanwhile we found a robust intracellular K^+^ decrease in macrophages treated with the K^+^ ionophore nigericin, we fail to detect a decrease of intracellular K^+^ when galvanic current was applicated (**Fig. S2B**). This data suggests that either a small and/or transient decrease of intracellular K^+^ could be induced by galvanic current or alternatively a dilution of intracellular K^+^ concentration should occur when galvanic current is applicated, and this could also explain the smaller concentration of IL-1β release induced by galvanic current compared to nigericin application (**Fig. 2C**). After galvanic current application we were able to detect the generation of the active p20 caspase-1 fragment, and processed IL-1β and GSDMD^NT^ (**Fig. 2D**). MCC950 was able to abrogate caspase-1 activation and the processed forms of IL-1β and GSDMD^NT^ (**Fig. 2D**), suggesting a functional caspase-1 activation and downstream signaling due to canonical NLRP3 activation and discarding the non-canonical NLRP3 activation that would result in GSDMD processing in the presence of MCC950.

**Figure 2.**
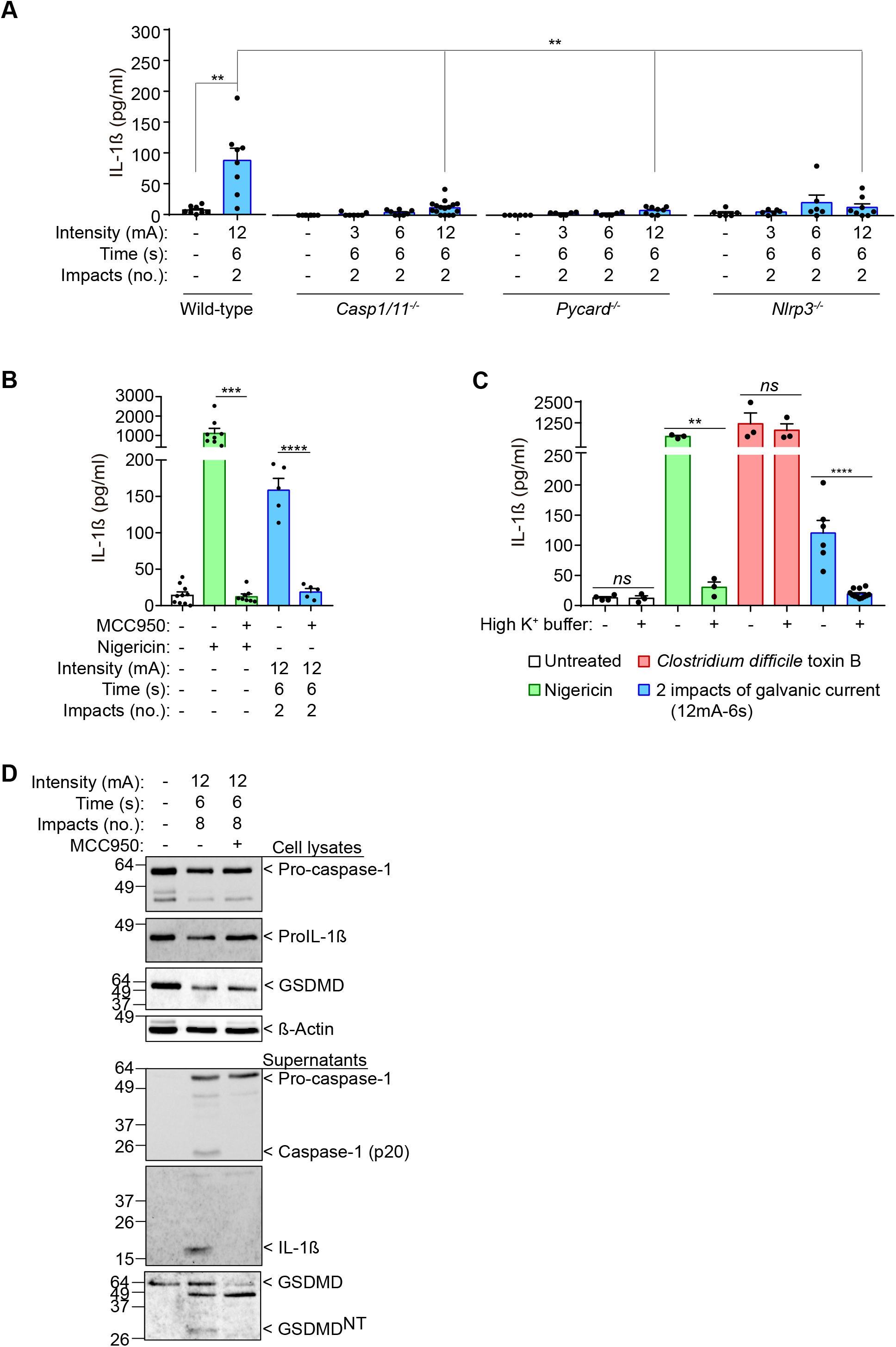
IL-1β release induced by galvanic current is dependent on the NLRP3 inflammasome. (**A**) IL-1β release from wild type, *Casp1/11*^-/-^, *Pycard*^-/-^ and *Nlrp3*^-/-^ mouse bone marrow derived macrophages (BMDMs) treated for 2 h with LPS (1 μg/ml) and then 2 impacts of different intensities of galvanic current (3, 6, 12 mA) for 6 seconds was applicated. Cells were then further cultured for 6 h before cytokine measurement in supernatant. Center values represent the mean and error bars represent s.e.m.; *n*= 6-16 samples of 10 independent experiments; LPS *vs* LPS+galvanic current in wild-type and wild-type *vs Nlrp3*^*-/-*^ unpaired *t*-test, wild-type *vs Casp1/11*^*-/-*^ and wild-type *vs Pycard*^*-/-*^ Mann-Whitney test, \*\**p*<0.005. (**B**) IL-1β release from wild type BMDMs treated as in A but applicating the NLRP3 specific inhibitor MCC950 (10 μM) 30 min before the galvanic current application and during the last 6 h of culture. As a control, cells were treated with nigericin (1,5 μM) instead galvanic current application. Center values represent the mean and error bars represent s.e.m.; *n*= 5-10 samples of 5 independent experiments; nigericin *vs* nigericin+MCC950 Mann-Whitney test, galvanic current *vs* galvanic current+MCC950 unpaired t-test, \*\*\*\**p*<0.0001 and \*\*\**p*<0.0005. (**C**) IL-1β release from wild type BMDMs treated as in A but applicating a buffer with 40 mM of KCl (high K^+^ buffer) during the last 6 h of culture. As controls, cells were treated with nigericin (1,5 μM) or *Clostridium difficile* toxin B (1 μg/ml) instead galvanic current application. Center values represent the mean and error bars represent s.e.m.; *n*= 3-12 samples of 4 independent experiments; unpaired *t*-test, \*\*\*\**p*<0.0001, \*\**p*<0.005 and *ns p*>0.05. (**D**) Immunoblot of cell extract and supernatants for caspase-1, IL-1β, GSDMD and β-actin from wild type BMDMs treated as in B, but with 8 impacts. Representative of *n*= 2 independent experiments.

### Galvanic current does not induce inflammasome-mediated pyroptosis

Since GSDMD was processed and the N-terminus detected upon galvanic current application, we next assessed pyroptosis by means of Yo-Pro-1 uptake to cells and LDH leakage from the cell. Two impacts of galvanic currents of different intensities (3, 6, 12 mA) for a period of 6 seconds (conditions that induce IL-1β release) were only inducing a significant, but slightly increase of cell death (**Fig. 3A**). This increase in cell death was not associated with the activation of the inflammasome, since it was also present in macrophages deficient on NLRP3, ASC or caspase-1/11 (**Fig. 3B**), suggesting that was independently of pyroptosis. Increasing the number or the time of 12 mA impacts applicated, resulted in a time-dependent increase of cell death (**Fig. 3A**), correlating with higher concentrations of IL-1β release (**Fig. 3C**). However, meanwhile IL-1β release was blocked by MCC950 (**Fig. 3C**), LDH release was not dependent on NLRP3 activation (**Fig. 3D**). This further corroborate that the NLRP3 activation is dependent on the intensity and time of galvanic current application. Similarly, two impacts of 12 mA for a period of 6 seconds were unable to induce plasma membrane permeabilization measured by Yo-Pro-1 uptake during a period of 3 h (**Fig. 3E**). Yo-Pro uptake increased over 3 h in an intensity dependent manner (3, 6, 12 mA) when 8 impacts were applicated during 6 seconds (**Fig. 3E**). This increase of plasma membrane permeabilization was not reverted after NLRP3 blocking with MCC950 or when ASC-deficient macrophages were used (**Fig. 3F**). All these results demonstrate that doses of galvanic current of 3 or 6 mA for impacts of 6 seconds do not compromise cell viability but are able to induce an inflammatory response dependent on NLRP3 activation, in contrast with current intensities of 12 mA that if prolonged in time could cause significant cell death independently of the inflammasome.

**Figure 3.**
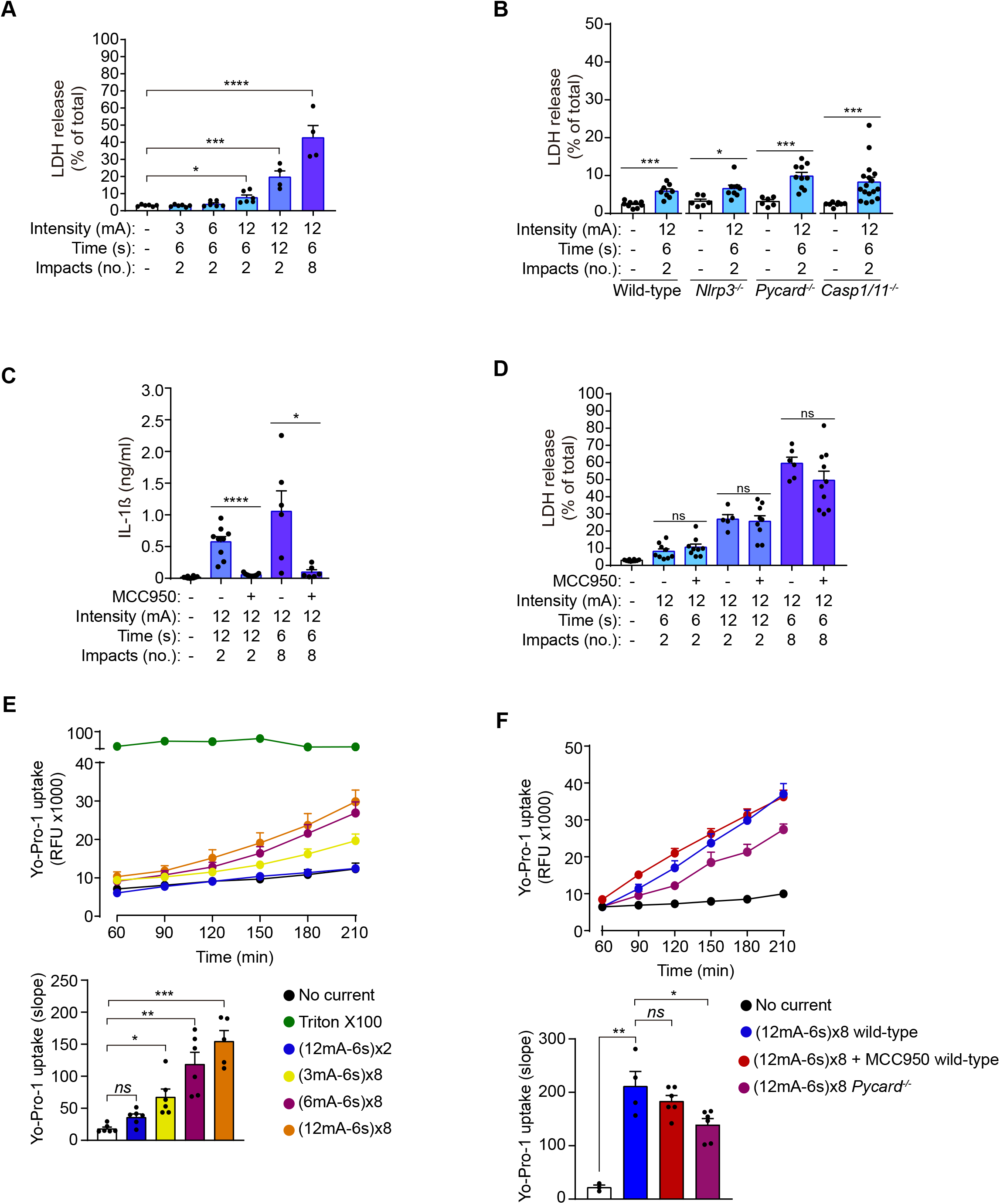
Galvanic current does not induce inflammasome-mediated pyroptosis. (**A**) Extracellular amount of LDH from mouse bone marrow derived macrophages (BMDMs) treated for 2 h with LPS (1 μg/ml) and then 2 or 8 impacts of different intensities of galvanic current (3, 6, 12 mA) for 6 or 12 seconds were applicated as indicated. Cells were then further cultured for 6 h before LDH determination in supernatant. Center values represent the mean and error bars represent s.e.m.; *n*= 3-4 samples of 6 independent experiments; Kruskal-Wallis test to compare LPS with increasing intensities of galvanic current, LPS *vs* LPS+ galvanic current (12mA-12s)x2 unpaired *t*-test, LPS *vs* LPS+ galvanic current (12mA-6s)x8, \*\*\*\**p*<0.0001, \*\*\**p*<0.0005 and \**p*<0.05. (**B**) Extracellular amount of LDH from wild type, *Nlrp3*^-/-^, *Pycard*^-/-^ and *Casp1/11*^-/-^ mouse BMDMs treated as in A. Center values represent the mean and error bars represent s.e.m.; *n*= 6-17 samples of 12 independent experiments; unpaired *t*-test except for *Casp1/11*^*-/-*^ comparison, \*\*\**p*<0.0005 and \**p*<0.05. (**C**) IL-1β release from wild type BMDMs treated as in A, but applicating the NLRP3 specific inhibitor MCC950 (10 μM) during the last 6 h of culture. Center values represent the mean and error bars represent s.e.m.; *n*= 6-10 samples of 5 independent experiments; unpaired *t*-test, \*\*\*\**p*<0.0001 and \**p*<0.005. (**D**) Extracellular amount of LDH from wild type BMDMs treated as in C. Center values represent the mean and error bars represent s.e.m.; *n*= 5-10 samples of 5 independent experiments; unpaired *t*-test, *ns p*>0.005. (**E**) Kinetic of Yo-Pro-1 uptake (upper panel) or slope of the uptake (lower panel) in wild type BMDMs treated for 2 h with LPS (1 μg/ml) and then with different intensities of galvanic current (as indicated) or with the detergent triton X-100 (1 %) during 3.5 h. Center values represent the mean and error bars represent s.e.m.; *n*= 3-6 of 3 independent experiments; Kruskal-Wallis test, \*\*\**p*<0.0005, \*\**p*<0.005 and *ns p*>0.05. (**F**) Kinetic of Yo-Pro-1 uptake (upper panel) or slope of the uptake (lower panel) in wild type or *Pycard*^-/-^ BMDMs treated as in E, but when indicated the NLRP3 specific inhibitor MCC950 (10 μM) was added after galvanic current application. Center values represent the mean and error bars represent s.e.m.; *n*= 3-6 samples of 3 independent experiments; unpaired *t*-test, \*\**p*<0.005, \**p*<0.05 and *ns p*>0.05.

### Galvanic current applicated in tendon increases inflammation *in vivo*

In order to study the effect of galvanic current *in vivo*, we found that application of 3 impacts of 3 mA of galvanic current during 3 seconds in the calcaneal tendon of mice resulted in an increase of the number of polymorphonuclear cells after 3 days when compared with tendons treated with needling alone (a puncture without current application, **Fig. 4A,B**). This increase returned to basal after 7 days and stayed low up to 21 days after galvanic current application (**Fig. 4B**). Similarly, the number of F4/80^+^ macrophages increased after 3 days of galvanic current application when compared to needling alone and returned to basal levels after 7 days (**Fig. 4C,D**). Other immune cell types detected in the tendon, as mastocytes, were not significantly increased by galvanic current application when compared to needling alone (**Fig. S3A**). Other histological features of the tendon (number of tenocytes, shape and area of tenocyte nuclei or neo-vascularization) were also not affected by the application of galvanic currents compared to needling alone (**Fig. S3B-E**).

**Figure 4.**
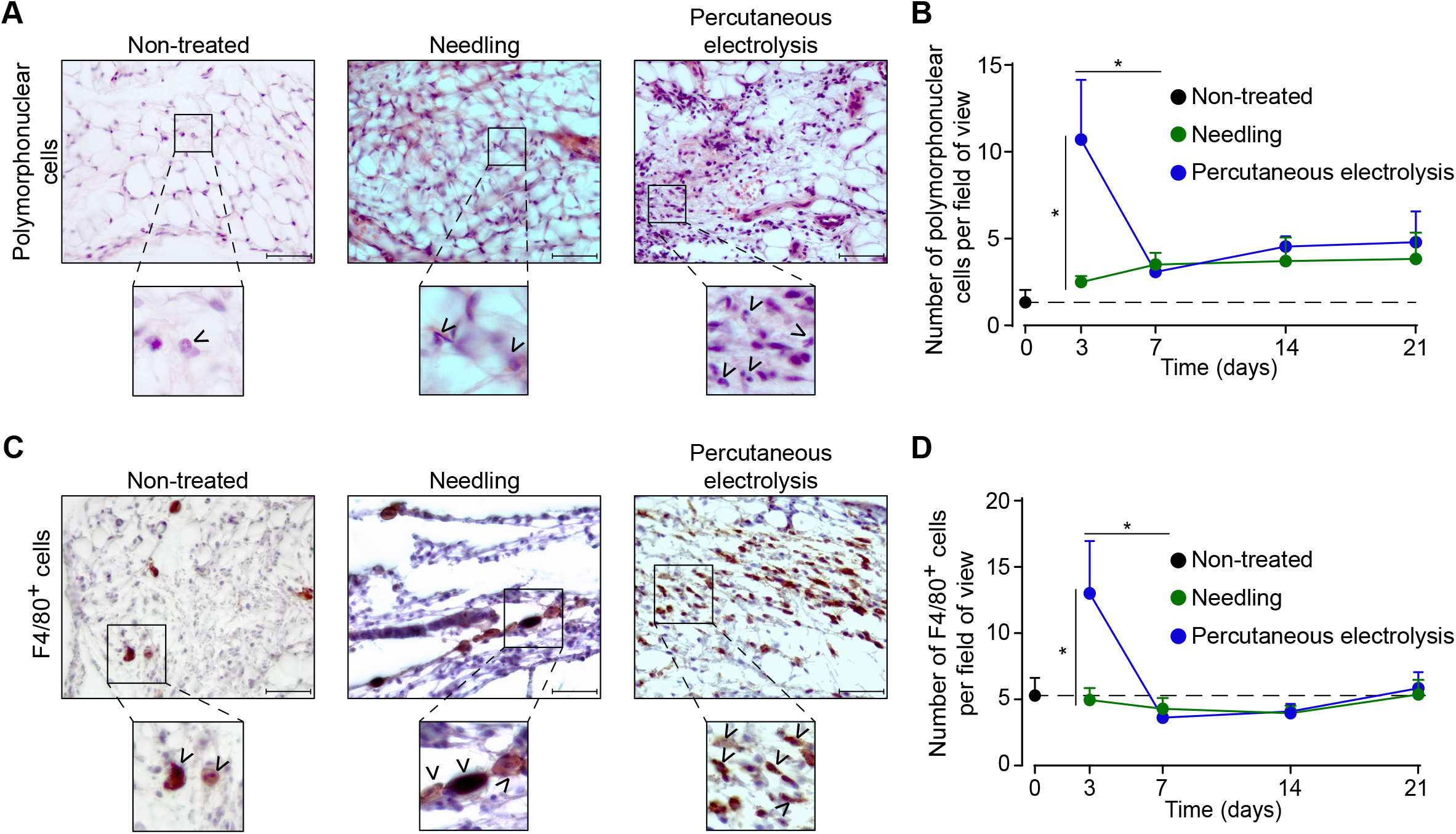
Galvanic current induces polymorphonuclear and macrophage infiltrate in the calcaneal tendon of mice. (**A**) Representative hematoxylin and eosin images of wild type mice calcaneal tendon after 3 days application of 3 punctures with needle (needling, green) or 3 impacts of 3 mA for 3 sec (blue). Scale bar: 50 μm. Magnification show the presence of polymorphonuclear cells (arrowheads). (**B**) Quantification of polymorphonuclear cells per field of view of calcaneal tendon sections treated and stained as described in A. Center values represent the mean and error bars represent s.e.m.; *n*= 7-8 independent animals; unpaired *t*-test, \**p*<0.005. (**C**) Representative immunostaining images for the macrophage marker F4/80 from the calcaneal tendon of wild type mice treated as described in A. Scale bar: 50 μm. Magnification show the presence of F4/80 positive cells (arrowheads). (**D**) Quantification of F4/80 positive cells per field of view of calcaneal tendon sections treated and stained as described in C. Center values represent the mean and error bars represent s.e.m.; *n*= 8 independent animals; Mann-Whitney test, \**p*<0.005.

We next assessed the expression of different pro-inflammatory cytokines in the calcaneal tendon after 3 days of 3 impacts of 3 mA of galvanic current application during 3 seconds. Expression of *Il6, Il1a* and *Il1b*, as well as the IL-1 receptor antagonist (*Il1rn*) and the chemokine *Cxcl10* were all increasing after percutaneous electrolysis when compared to needling alone (**Fig. 5A**). Different NLRP3 inflammasome genes also exhibit an increase in expression (*Nlrp3, Pycard, Casp1*) when galvanic current was applicated, but this increase was not significantly when compared to needling (**Fig. 5B**). *Gsdmd* expression was not upregulated in the tendons after galvanic current application (**Fig. 5B**). These data suggest that galvanic current induces an inflammatory response driven by the infiltration of polymorphonuclear cells and macrophages, together an increase of the expression of several cytokines and chemokines.

**Figure 5.**
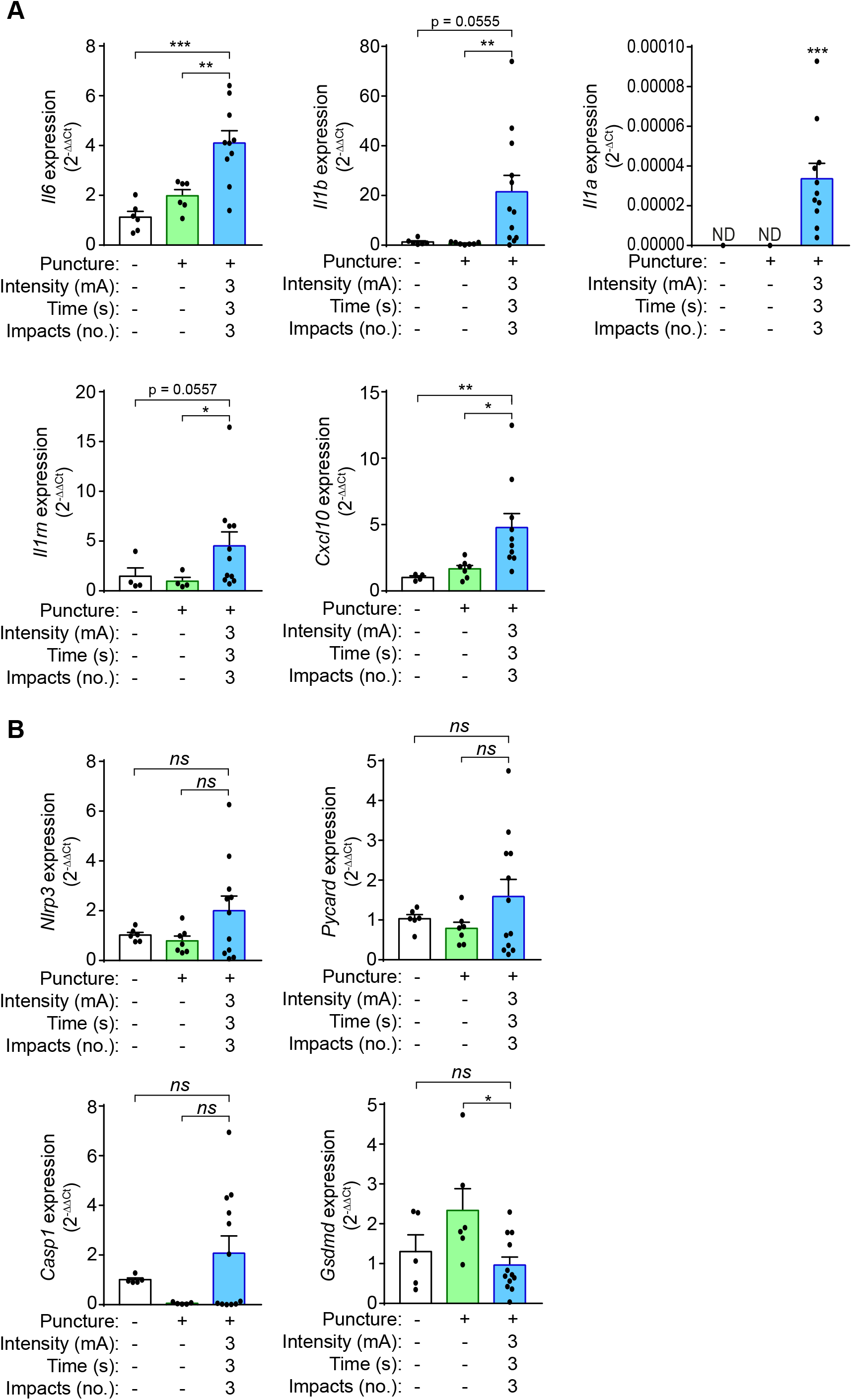
Galvanic current induces proinflammatory cytokine expression in the calcaneal tendon of mice. (**A**,**B**) Quantitative PCR for the indicated genes normalized to *Actb* in the calcaneal tendon of wild type mice after 3 days application of 3 punctures with needle (needling, green) or 3 impacts of 3 mA for 3 sec (blue), and compared to the expression of genes in non-treated tendons. Center values represent the mean and error bars represent s.e.m.; *n*= 4-12 independent animals; for *Il6, Nlrp3, Pycard* and *Gsdmd* unpaired *t*-test, for *Il1b* untreated *vs* galvanic current Mann-Whitney and puncture *vs* galvanic current unpaired *t*-test, for *Cxcl10, Il1rn* and *Casp1/11* Mann-Whitney test, for *Il1a* one sample Wilcoxon test (ND: non detected), \*\*\**p*<0.0005, \*\**p*<0.005, \**p*<0.05 and *ns p*>0.05.

### The NLRP3 inflammasome controls the *in vivo* inflammatory response induced by galvanic current

In order to evaluate if the NLRP3 inflammasome mediates the inflammatory response in tendons after percutaneous electrolysis, we applied galvanic currents in the calcaneal tendon of *Nlrp3*^-/-^ mice. Application of 3 impacts of 3 mA of galvanic current for 3 seconds in the calcaneal tendon of *Nlrp3*^-/-^ mice resulted in a significant reduction of *Il1b, Il1rn* and *Cxcl10* expression after 3 days when compared to wild-type mice (**Fig. 6A**). Specific inflammasome associated genes, as *Pycard, Casp1* or *Gsdmd* (except for *Nlrp3*) where not affecting their expression in the calcaneal tendon of *Nlrp3*^-/-^ mice after 3 days of galvanic current application when compared to wild type mice (**Fig. 6B**). Surprisingly, galvanic current produced a tendency to increase the expression of *Il6* in the tendons of *Nlrp3*^-/-^ after 3 days (**Fig. 6C**) and in parallel, the number of polymorphonuclear cells was also increased (**Fig. 6D**). However, the number of macrophages was not affected in the *Nlrp3*^-/-^ calcaneal tendon when galvanic current was applicated (**Fig. 6D**). We also confirmed a decrease of *Il1b* and *Cxcl10* expression in the tendons of *Pycard*^-/-^ mice after 3 days of galvanic current application (**Fig. S4**), suggesting that the NLRP3 inflammasome is important to modulate part of the inflammatory response after galvanic current application.

**Figure 6.**
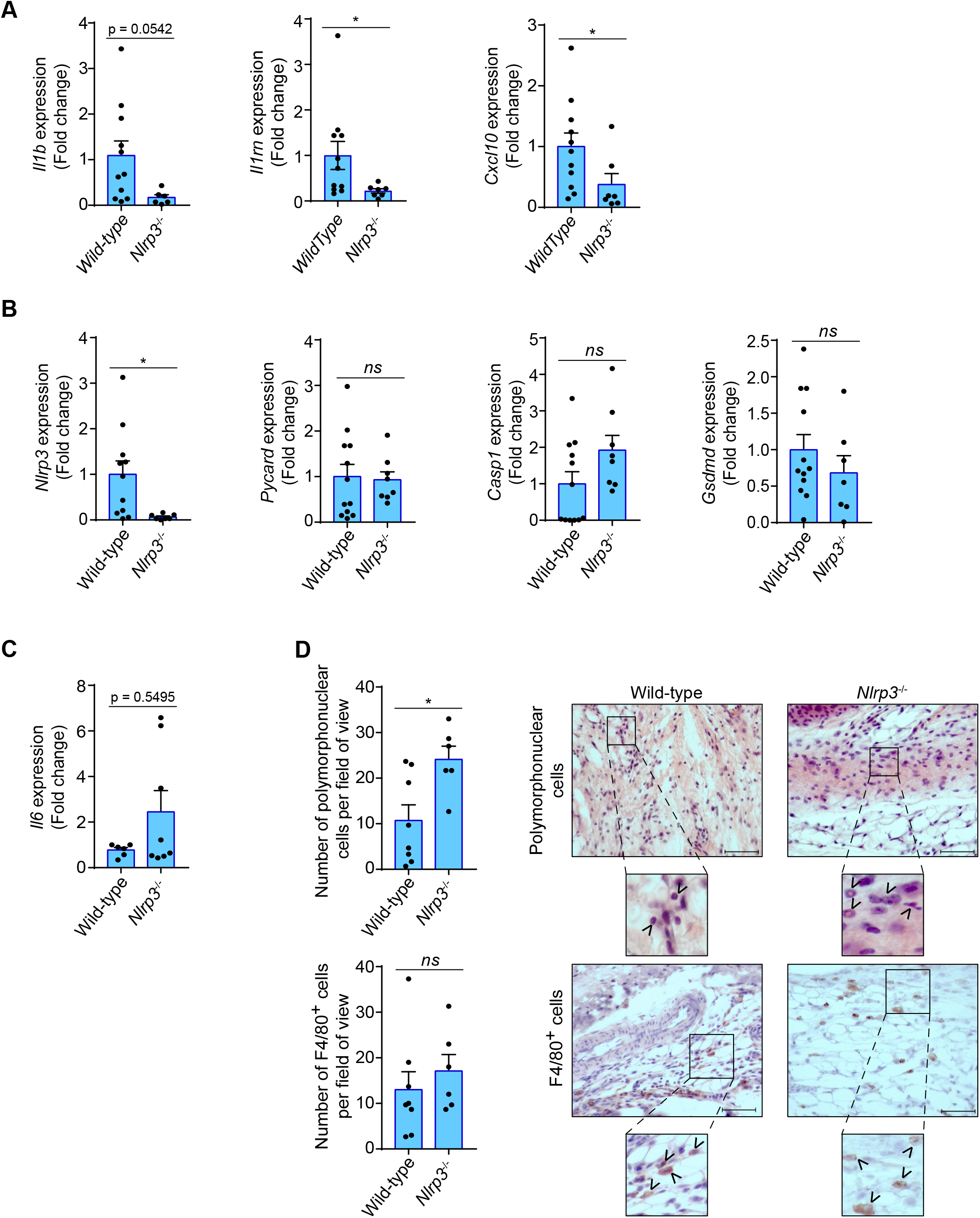
Inflammatory response in the calcaneal tendon of *Nlrp3*^-/-^ mice after galvanic current application. (**A**-**C**) Quantitative PCR for the indicated genes in the calcaneal tendons of *Nlrp3*^-/-^ mice (calculated as 2^-ΔΔCt^) normalized to the expression in wild type (calculated as 2^-ΔΔCt^) after 3 days application of 3 impacts of 3 mA for 3 sec. Center values represent the mean and error bars represent s.e.m.; *n*= 3-12 independent animals; for *Il1b, Nlrp3, Pycard, Casp1/11* and *Gsdmd* unpaired *t*-test, for *Il1rn* and *Cxcl10* Mann-whitney test, \**p*<0.05 and *ns p*>0.05. (**D**) Quantification of polymorphonuclear (top) and F4/80 positive cells (bottom) per field of view from wild type and *Nlrp3*^-/-^ mice calcaneal tendon treated as in A. Center values represent the mean and error bars represent s.e.m.; *n*= 6-8 independent animals; unpaired *t*-test, \**p*<0.05 and *ns p*>0.05. representative hematoxylin and eosin images (top) and F4/80 immunostaining (bottom) of calcaneal tendon quantified. Scale bar: 50 μm. Magnification show the presence of polymorphonuclear (top) or F4/80 cells (bottom) denoted by arrowheads.

### The NLRP3 inflammasome induces a tissue regenerative response to galvanic current application

Galvanic current application has been widely used to resolve chronic tendinopathies (Abat et al., 2016; Rodríguez-Huguet et al., 2020; Valera-Garrido et al., 2014), and here we present the case of a 6 weeks resolution of lateral epicondylitis after four sessions of percutaneous electrolysis with an intensity of 3 mA of galvanic current application for 3 sec, 3 times (3:3:3) (**Fig. 7A**), according to the protocol previously described by Valera-Garrido and Minaya-Muñoz (Valera-Garrido and Minaya-Muñoz, 2016). During this tissue regeneration, the production of new extracellular matrix is a key process (Shook et al., 2018; Wynn, 2008). In order to investigate if the inflammatory response mediated by the NLRP3 inflammasome after galvanic current application is important for tissue regeneration, we measured *Tgfb1* expression as a key factor inducing collagen production. We found that *in vivo* the expression of *Tgfb1* after 3 days of galvanic current application in the calcaneal tendon of mice was dependent on NLRP3 (**Fig. 7B**). In line, after 7 days of percutaneous electrolysis the levels of type III collagen were decreased, with a parallel increase of type I collagen when compared to needling alone (**Fig. 7C, S5E**). However, percutaneous electrolysis did not affect collagen fiber properties measured (width, length, strength or angle) when compared to needling alone (**Fig. S5A-D**). The increase of type I collagen after 7 days of galvanic current application was reduced in *Nlrp3*^-/-^ mice (**Fig. 7D**), suggesting that the NLRP3 inflammasome could control the response of galvanic current inducing type I collagen. Overall, we found that galvanic current application is able to activate the NLRP3 inflammasome and induce the release of IL-1β, initiating an inflammatory response that could lead to the regeneration of the tendon (**Fig. S6**).

**Figure 7.**
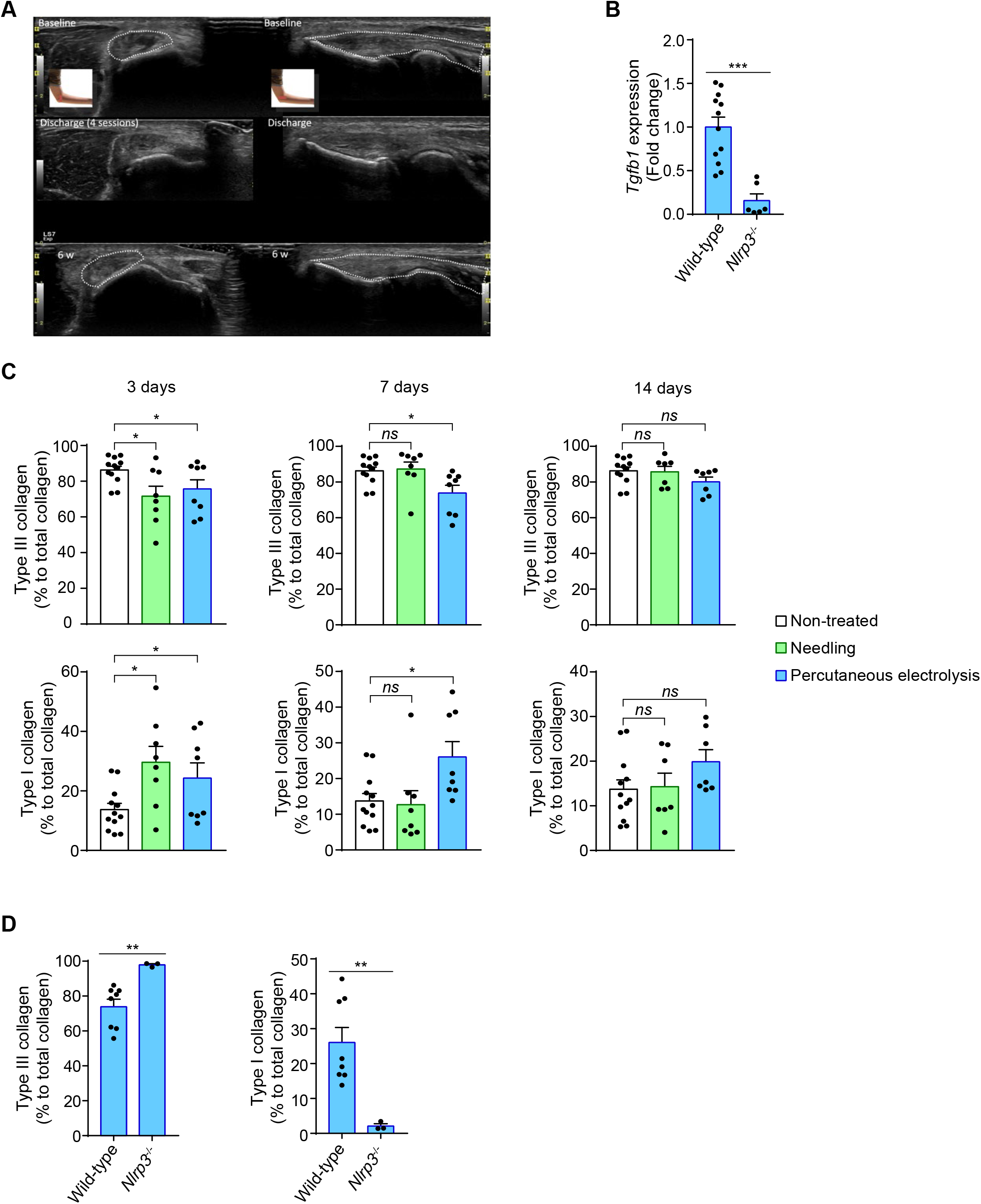
Galvanic current increase of type I collagen via NLRP3 inflammasome. (**A**) Ultrasound scanning of the right elbow of a patient of with lateral epicondylalgia for 6 months of evolution, with pain an functional impairment. Baseline represent the image before galvanic current application. Discharge, represent the image after 4 sessions of percutaneous electrolysis with an intensity of 3 mA of galvanic current application for 3 sec, 3 times (3:3:3). 6 w, represent the image after 6 weeks of the last session of percutaneous electrolysis. (**B**) Quantitative PCR for *Tgfb1* in the calcaneal tendons of *Nlrp3*^-/-^ mice (calculated as 2^-ΔΔCt^) normalized to the expression in wild type (calculated as 2^-ΔΔCt^) after 3 days application of 3 impacts of 3 mA for 3 sec. Center values represent the mean and error bars represent s.e.m.; *n*= 6-12 independent animals; Mann-Whitney test, \*\*\**p*<0.0005. (**C**,**D**) Quantification of the collagen type I and III in calcaneal tendon sections stained with picrosirius red from wild type (C,D) and *Nlrp3*^-/-^ (D) mice after 3, 7 or 14 days (C) or 3 days (D) application of punctures with needle (needling, green) or 3 impacts of 3 mA for 3 sec (blue), or in treated tendons (white). Center values represent the mean and error bars represent s.e.m.; *n*= 3-12 independent animals; for 3 days and panel D unpaired *t*-test, for 7 days non-treated *vs* needling Mann-Whitney test and non-treated *vs* percutaneous electrolysis unpaired *t*-test, for 14 days untreated *vs* needling unpaired *t*-test and untreated *vs* percutaneous electrolysis Mann-Whitney test, \*\**p*<0.005, \**p*<0.05 and *ns p*>0.05.

## DISCUSSION

In this study we demonstrate how galvanic current application induces in macrophages a pro-inflammatory signature, mainly characterized by the activation of the NLRP3 inflammasome and the release of mature IL-1β. Inflammation is an important initial response that promotes tissue repair and wound healing, however excessive inflammation could lead to chronic inflammation and fibrosis (Alegre et al., 2017; Eming et al., 2017; Gaul et al., 2020). The NLRP3 inflammasome is a key pathway to control inflammation in the absence of pathogenic microorganisms (in sterile conditions) and its stimulation favors the activation of caspase-1 and the processing of GSDMD, that execute a pro-inflammatory type of cell death called pyroptosis (Broz et al., 2020; Broz and Dixit, 2016; Liston and Masters, 2017). Pyroptosis compromises the integrity of the cellular plasma membrane and results in the uncontrolled release of intracellular content, including the release of the potent pro-inflammatory cytokines IL-1β and IL-18 (Broz et al., 2020). Pyroptosis also leads to the release of inflammasome oligomers that spread pro-inflammatory signaling and drives fibrosis (Baroja-Mazo et al., 2014; Franklin et al., 2014; Gaul et al., 2020). Here we found that galvanic current application, a technique that has been widely used to resolve chronic lesions in clinic (Valera-Garrido et al., 2014), was able to activate the NLRP3 inflammasome and induce IL-1β release, but with very little associated pyroptotic cell death. This could be due to two potentially different mechanisms: (*i*) the finding of an alternative processing of GSDMD after galvanic current application that was independent on NLRP3 and could inactivate its N-terminal lytic domain, as has been previously found for caspase-3 processing GSDMD (Taabazuing et al., 2017); and/or (*ii*) the small amounts of GSDMD^NT^ found that could result in a small number of pores at the plasma membrane and facilitates their repair by the endosomal sorting complexes required for transport machinery leading to an hyperactive state of the macrophage (Evavold et al., 2018; Rühl et al., 2018). During this state of the macrophage, IL-1β is released in the absence of cell death (Evavold et al., 2018). However, an increase of the intensity and the time of galvanic current application leads to an increase in cell death, that was independent on the inflammasome and could be related to the technique *per se*. Therefore, clinical application of current intensities above 6 mA would probably lead to necrosis of the tissue and not to an efficient reparative process. Galvanic currents of 3 and 6 mA application for 2 impacts of 6 seconds are both able to induce NLRP3 inflammasome activation *in vitro* and also lead to phenotypic changes in the tendon *in vivo*. This is in line with the fact that 3 mA of galvanic current are able to induce a clinically relevant regeneration of lesions (García Vidal et al., 2019; Margalef et al., 2019; Medina i Mirapeix et al., 2019; Valera-Garrido et al., 2014). High intensity doses for long periods of time or repeated impacts could induce massive tissue necrosis and therefore are not recommended in the clinical practice.

The activation of the NLRP3 inflammasome induced by galvanic currents was found dependent on K^+^ efflux, as extracellular high concentrations of K^+^ was able to block IL-1β release, but surprisingly galvanic current application was not resulting in a detectable intracellular K^+^ decrease. This oppose the effect of the well-studied K^+^ ionophore nigericin that was able to induce a dramatic decrease of intracellular K^+^ in accordance to previous publications (Muñoz-Planillo et al., 2013; Petrilli et al., 2007; Próchnicki et al., 2016). It might be that galvanic currents would induce a slight decrease of intracellular K^+^ not detectable by the technique used in this study, but enough to result in NLRP3 activation. In fact, the amount of IL-1β released from galvanic current activated macrophages was lower than when macrophages were activated with nigericin, denoting a correlation to the decrease in intracellular K^+^. The low NLRP3 activation induced by galvanic current application could result in a moderate inflammatory response *in vivo* beneficial for tissue regeneration. In fact, NLRP3 was important to induce an inflammatory response *in vivo* with elevation of different cytokines including *Il1b* or *Cxcl10*, but contrary affecting *Il6* production and the deficiency of NLRP3 leads to an increase of polymorphonuclear cells. Also, exacerbated NLRP3 activation could led to fibrosis (Alegre et al., 2017; Gaul et al., 2020), denoting that NLRP3 could control collagen deposition. The mild activation of NLRP3 found after galvanic current application was associated to increase production of *Tgfb1* and an increase of collagen type I vs type III in tendonds. This could explain the beneficial regenerative response of the application of galvanic current in tendon lesions (Abat et al., 2016; Rodríguez-Huguet et al., 2020; Valera-Garrido et al., 2014, 2013).

Therefore, this study reports how galvanic current is a feasible technique applicated *in vivo* to activate the NLRP3 inflammasome and induce a local inflammatory response to enhance a collagen-mediated regeneration process in the tendon, establishing the molecular mechanism of percutaneous electrolysis for the treatment of chronic lesions.

## MATERIAL AND METHODS

### Animals and percutaneous needle puncture procedure

All experimental protocols for animal handling were refined and approved by the local animal research ethical committee (references 241/2016 and 541/2019) and Animal Health Service of the General Directorate of Fishing and Farming of the Council of Murcia (*Servicio de Sanidad Animal, Dirección General de Ganadería y Pesca, Consejería de Agricultura y Agua de la Región de Murcia*, reference A13160702). C57/BL6 mice (wild-type) were obtained from the Jackson Laboratories. NLRP3-deficient mice (*Nlrp3*^*-/-*^) and Caspase-1/11-deficient mice (*Casp-1/11*^*-/-*^) in C57/BL6 background were a generous gift of I. Coullin. For all experiments, mice between 8-10 weeks of age were used. Mice were bred in specific pathogen-free conditions with 12:12 h light-dark cycle and used in accordance with the *Hospital Clínico Universitario Vírgen de la Arrixaca* animal experimentation guidelines, and the Spanish national (RD 1201/2005 and Law 32/2007) and EU (86/609/EEC and 2010/63/EU) legislation. Percutaneous needle puncture was performed with 16G and 13 mm needles (Agupunt) in the calcaneal tendon in isoflurane (Zoetis) anesthetized mice; galvanic current was applicated using Physio Invasiva® equipment (Prim) delivering three impacts of 3 mA for 3 seconds and compared to a puncture without current application. Paws without puncture were also used as controls. 3, 7, 14 and 21 days after puncture, animals were euthanized and paws were collected for histopathology or gene expression. Only calcaneal tendon was dissected for gene expression and the zone between gastrocnemius and calcaneus, including tendon, adipose tissue, tibia and peroneus, was dissected for histopathology.

### Patient

A male patient of 36 year old with lateral epicondylalgia in the right elbow for 6 months of evolution, with pain an functional impairment. Resistant to conventional treatments (physiotherapy, oral non-steroidal anti-inflammatory and local corticoid infiltrations). Ultrasound analysis show extensor joint tendon degeneration correlating with positive orthopedic tests. The patient was subjected to four sessions of percutaneous electrolysis with an intensity of 3 mA of galvanic current application for 3 sec, 3 times (3:3:3), according to the protocol by Valera-Garrido and Minaya-Muñoz (Valera-Garrido and Minaya-Muñoz, 2016).

### Cell culture and treatments

Bone marrow-derived macrophages (BMDMs) were obtained from wild-type, *Casp1/11*^-/-^, *Nlrp3*^-/-^ and *Pycard*^-/-^ mice. Cells were differentiating for 7 days in DMEM (Lonza) supplemented with 25% of L929 medium, 15% fetal bovine serum (FCS, Life Technologies), 100 U/ml penicillin/streptomycin (Lonza), and 1% L-glutamine (Lonza). After differentiation, cells were primed for 2 h with 1 µg/ml *E. coli* lipopolysaccharide (LPS) serotype O55:B5 at (Sigma-Aldrich) or for 4 h with 20 ng/ml recombinant mouse IL-4 (BD Pharmigen). Cells were then washed twice with isotonic buffer composed of 147 mM NaCl, 10 mM HEPES, 13 mM glucose, 2 mM CaCl_2_, 1 mM MgCl_2_, and 2 mM KCl, pH 7.4, and then treated in OptiMEM (Lonza) with different intensities and time of galvanic current (as indicated in the text and figure legends) using an ad hoc adaptor for 6 well plates (**Fig. S1**), and then cultured for 6 h. Alternatively and as a positive control, after LPS-priming macrophages were treated for 6 h in OptiMEM with 1.5 µM nigericin (Sigma-Aldrich) or 1 µg/ml *Clostridium difficile* toxin B (Enzo Life Sciences) to activate NLRP3 and Pyrin inflammasomes respectively. In some experiments, cells were treated with 10 µM of the NLRP3 inflammasome inhibitor MCC950 (CP-456773, Sigma-Aldrich) after LPS priming and during inflammasome activation.

### LDH release, Yo-Pro uptake assay and K^+^ measurements

The presence of lactate dehydrogenase (LDH) in cell culture supernatants was measured using the Cytotoxicity Detection kit (Roche), following manufacturer’s instructions. It was expressed as the percentage of the total amount of LDH present in the cells. For Yo-Pro uptake, macrophages were preincubated for 5 min at 37 °C with 2.5 µM of Yo-Pro-1 iodide (Life Technologies) after galvanic current application or 1% triton X100 (Sigma-Aldrich) application. Yo-Pro-1 fluorescence was measured after the treatments every 5 minutes for the first 30 min and then every 30 min for the following 3 h with an excitation wavelength of 478 ± 20 nm and emission of 519 ± 20nm in a Synergy neo2 multi-mode plate reader (BioTek). Intracellular K^+^ was quantified from macrophages lysates as already reported (Compan et al., 2012) and measured by indirect potentiometry on a Cobas 6000 with ISE module (Roche).

### Western blot and ELISA

After cell stimulation, cells extracts were prepared in cold lysis buffer and incubated at 4°C for 30 min and then centrifuged at 12856 x*g* for 10 min at 4°C. Cells supernatants were centrifuged at 12856 x*g* for 30 seconds at 4°C and concentrated by centrifugation at 11000 x*g* for 30 min at 4°C through a column with a 10 kDa cut-off (Merk-Millipore). Cell lysates and concentrated supernatants were mixed with loading buffer (Sigma), boiled at 95°C for 5 min, resolved in 15% polyacrylamide gels and transferred to nitrocellulose membranes (BioRad). Different primary antibodies were used for the detection of interest proteins: anti-IL-1β rabbit polyclonal (1:1000, H-153, SC-7884, Santa Cruz), anti-caspase-1 (p20) mouse monoclonal (1:1000, casper-1, AG-20B-0042, Adipogen), anti-gasdermin D rabbit monoclonal (1:2000, EPR19828, ab209845, Abcam) and anti-β-Actin mouse monoclonal (1:10000, Santa Cruz). Appropriate secondary antibody conjugated with HRP was used at 1:5000 dilution (Sigma) and developed with ECL plus (Amhershan Biosciences) in a ChemiDoc HDR (BioRad). Uncropped Western blots are shown in **Figure 2-source data 1 and 2**. The concentration of IL-1β, TNF-α and IL-6 in cell supernatants was determined by ELISA following the manufacturer’s instructions (R&D Systems). Results were read in a Synergy Mx plate reader (BioTek).

### Quantitative reverse transcriptase-polymerase chain reaction (RT-PCR) analysis

Total RNA extraction was performed using macrophages or mice tendons dissected as described above. Macrophage total RNA extraction was performed using the RNeasy Mini Kit (Qiagen) following manufacturer’s instructions. Total RNA extraction from mice tendons was performed using Qiazol lysis reagent (Qiagen) and samples were homogenized using an Omni THQ homogenizer. After homogenization, samples were incubated 5 min at room temperature and centrifuged at 12000 *xg* for 15 min at 4°C. After centrifugation, upper phase was collected and one volume of 70% ethanol was added. Samples were loaded in RNeasey Mini Kit columns and total RNA isolation was performed following manufacturer’s instructions. In both cases a step with a treatment with 10 U/μl DNase I (Qiagen) was added during 30 min. Reverse transcription was performed using iScript cDNA Synthesis kit (BioRad). The mix SYBR Green Premix ExTaq (Takara) was used for quantitative PCR in an iCyclerMyiQ thermocycler (BioRad). Specific primers were purchased from Sigma (KiCqStart SYBR Green Primers) for the detection of the different genes. Relative expression of genes was normalized to the housekeeping gene *Actb* using the 2^-ΔCt^ method and for the expression in tendon then normalized to mean Ct value of non-treated samples using the 2^-ΔΔCt^ method (value shown in figures). When expression in non-treated samples was below threshold and was non detected (ND), 2^-ΔCt^ values are shown in the figures. To compare gene expression between wild-type and knock-out mice, the fold change of 2^-ΔΔCt^ values of the knock-out mice was calculated respect the average of the 2^-ΔΔCt^ values of the wild-type mice.

### Histopathology

Mice paws were fixed using 4% *p*-formaldehyde (Sigma-Aldrich) for at least 24 h, processed, paraffin-embedded, and sectioned in 4 μm slides. Hematoxylin and eosin stained slices were initially evaluated in a 0 to 3 qualitative scale, being 0 control (healthy tendon) conditions, 1 mild, 2 medium, and 3 severe, for inflammatory infiltrate and tendon cellularity grade (the median value for each of the paws was used as the final value represented in the figures), the number of polymorphonuclear cells was quantified by counting 3 different fields for each sample, attending to nuclear morphology of the cells in adipose tissue next to calcaneal tendon, the number and area of tenocytes nuclei was evaluated using the FIJI macro based on a manual threshold to select nuclei and evaluating the different parameters measured with the “Analyze particles” tool. Sirius red staining was performed in the slides using the Picro Sirius red stain kit (Abcam) following the manufacturer’s instructions and polarized light pictures (**Fig. S5E**) were used to quantify the type of collagen by converting pictures to SHG color and then using the CT-Fire algorithm to calculate width, length, straightness and angle of collagen fibers in these pictures (Liu et al., 2017). Immunohistochemistry with anti-F4/80 rat monoclonal antibody (MCA497GA, BioRad) was used for the quantification of macrophages by counting 3 different fields for each sample, attending to stained cells in adipose tissue next to calcaneal tendon. All slides were examined with a Zeiss Axio Scope AX10 microscope with 20x and 40x objectives (Carl Zeiss) and pictures were taken with an AxioCam 506 Color (Carl Zeiss).

### Statistics

Statistical analyses were performed using GraphPad Prism 7 (Graph-Pad Software, Inc). A Shapiro-Wilk normality test was initially performed to all groups to decide the analysis type to be used. For two-group comparisons, nonparametric Mann-Whitney *U* test (without making the assumption that values are normally distributed) or the parametric unpaired *t*-test (for normal distributed data) were used to determine the statistical significance. For more than two group comparisons, one-way ANOVA test (for normal distributed data) or nonparametric Krustal-Wallis test (without making the assumption that values are normally distributed) were used to determine the statistical significance. Data are shown as mean values and error bars represent standard error from the number of independent assays indicated in the figure legend, which are also overlaid in the histograms as dot-plotting. *p* value is indicated as \**p* <0.05; \*\**p* <0.01; \*\*\**p* <0.001; \*\*\*\**p* <0.0001; p >0.05 not significant (*ns*).

## Acknowledgments

We thanks M.C. Baños (IMIB-Arrixaca, Murcia, Spain) for technical assistance with molecular and cellular biology, Antonio García Martínez (CESMAR Electromedicina) for electrode development for *in vitro* application of galvanic current, Darío Peñín Franch for helping in the development of the plugin to measure different types of collagen and F. Noguera and M. Martínez (IMIB-Arrixaca, Murcia, Spain) for running Hitachi ion detection system and the members of the Pelegrin’s laboratory for comments and suggestions thought the development of this project. We also want to acknowledge the support of the SPF-animal house from IMIB-Arrixaca.

## Funding

A.P-F. was supported by MVClinic and Prim. This work was supported by grants to P.P. from *FEDER/Ministerio de Ciencia, Innovación y Universidades – Agencia Estatal de Investigación* (grant SAF2017-88276-R and PID2020-116709RB-I00), *Fundación Séneca* (grants 20859/PI/18 and 21081/PDC/19), and the European Research Council (ERC-2013-CoG grant 614578 and ERC-2019-PoC grant 899636).

## Author contributions

A.P-F., performed all the experimental work; A.P-F., J.A.G-V., C.M.M., P.E-R. performed *in vivo* animal manipulation and histology; A.P-F. and P.P. analyzed the data, interpreted results, conceived the experiments, prepared the figures and paper writing; F.V-G. and F.Mi-M. provided clinical data and conceptual supervision of the study. F.Me-M. and P.P. conceived the project, provided funding and overall supervision of this study.

## Declaration of interests

F.Mi.-M. and F.V.-G. are employees of MVClinic Institute. A.P-F. contract was supported by MVClinic Institute and Prim. P.P. declares that he is an inventor in a patent filled on March 2020 by the Fundación para la Formación e Investigación Sanitaria de la Región de Murcia (PCT/EP2020/056729) for a method to identify NLRP3-immunocompromised sepsis patients. P.P. is consultant of Glenmark Pharmaceutical. The remaining authors declare no competing interests.

